# ZapA stabilizes FtsZ filament bundles without slowing down treadmilling dynamics

**DOI:** 10.1101/580944

**Authors:** Paulo Caldas, Mar López-Pelegrín, Daniel J.G. Pearce, Nazmi B. Budanur, Jan Brugués, Martin Loose

## Abstract

For bacterial cell division, treadmilling filaments of FtsZ organize into a ring-like structure at the center of the cell. What governs the architecture and stability of this dynamic Z-ring is currently unknown, but FtsZ-associated proteins have been suggested to play an important role. Here, we used an *in vitro* reconstitution approach combined with fluorescence microscopy to study the influence of the well-conserved protein ZapA on the organization and dynamics of FtsZ filaments recruited to a supported membrane. We found that ZapA increases the spatial order and stabilizes the steady-state architecture of the FtsZ filament network in a highly cooperative manner. Despite its strong influence on their large-scale organization, ZapA binds only transiently to FtsZ filaments and has no effect on their treadmilling velocity. Together, our data explains how FtsZ-associated proteins can contribute to the precision and stability of the Z-ring without compromising treadmilling dynamics.

## Introduction

For cytokinesis, bacteria need to build two new cell poles at the division site. This process is initiated by a dynamic ring of FtsZ filaments, which forms at the mid-cell early during the cell cycle, where it persists during the inward growth of the cell septum until it disassembles just before division is completed^1-4^. Recent studies have shown that this Z-ring not only defines the location of divisome assembly, but that FtsZ treadmilling drives the movement of peptidoglycan synthases around the cell diameter, which in turn controls remodeling of the cell wall at the division site^5,6^. Due to the important role of FtsZ polymerization dynamics for cell wall synthesis, the spatial and temporal organization of filaments within the Z-ring needs to be tightly controlled. Indeed, bacteria contain various proteins that directly bind to FtsZ and contribute to the precision of cell division. For example, mutants of *Escherichia coli* lacking one of the FtsZ-associated proteins ZapA, ZapB, ZapC or ZapD are usually longer than wild-type cells with a more heterogeneous cell length distribution. Furthermore, they show mislocalized or misaligned Z-rings and skewed division septae^7-10^. Despite not being structurally related, all these proteins have similar functions *in vivo* with their corresponding phenotype becoming even more severe in cells missing more than one Zap protein. However, the mechanism by which FtsZ-associated proteins contribute to the stability of the Z-ring and precision of cell division is currently not known.

The best characterized of those proteins is ZapA, a widely conserved protein critical for the positioning and stability of the division machinery^7,8,11^. Structural studies on ZapA from *E. coli* and *Pseudomonas aeruginosa* revealed that ZapA has two distinct domains. A globular N-terminal portion that forms the FtsZ-binding domain, and a C-terminal part that forms a 14-turn alpha helix, responsible to promote the oligomerization of the protein into a pseudosymmetric tetramer^12-14^. *In vitro* experiments using purified proteins showed that ZapA promotes the formation of stable FtsZ protofilament bundles and reduces the GTPase activity of FtsZ in solution^15-17^. These two properties, however, would potentially interfere with the function of FtsZ treadmilling to distribute cell wall synthases. Accordingly, it is currently not clear how ZapA can contribute to the stability of the cell division machinery without having a negative effect on the polymerization dynamics of its main organizer, FtsZ.

Here, we used an *in vitro* reconstitution approach recapitulating Z-ring assembly and early events during divisome maturation. Our biomimetic system combines purified proteins, supported bilayers and total internal reflection fluorescence (TIRF) microscopy. This allowed us to directly visualize the polymerization of FtsZ with its membrane anchor FtsA into filament networks and the influence of ZapA on their architecture and dynamics.

Using a novel automated image analysis, we studied the behavior of FtsZ and ZapA on three different spatial scales: First, we analyzed the spatiotemporal pattern of the membrane-bound filament network; next, we studied the underlying polymerization dynamics of filament bundles, and finally, we quantified the behavior of single molecules. Our experiments and quantitative analyses reveal that ZapA is able to align FtsZ filaments in a highly cooperative manner, giving rise to two distinct states: a state of low spatial order with fast reorganization dynamics and a state of high spatial order and slow reorganization dynamics. Importantly, we show that the treadmilling velocities are identical in these two regimes, demonstrating that ZapA is able to stabilize filament bundles without affecting FtsZ polymerization dynamics. Our results suggest that the function of the ZapA-FtsZ interaction is to cooperatively align FtsZ filaments at the division site, which defines the track for treadmilling and thereby increases the spatiotemporal precision of cell division.

## Results

### ZapA increases the bundle width of membrane-bound FtsZ filaments

To study the effect of ZapA on the architecture and dynamics of the FtsZ filament network, we took advantage of an *in vitro* system based on supported lipid bilayers and purified proteins (Fig. 1A). As reported previously, FtsZ and FtsA form treadmilling filaments on supported membranes when incubated in the presence of ATP and GTP and at a concentration ratio similar to that found in the living cell (FtsZ : FtsA = 1.5 μM : 0.5 μM). These filaments further self-organize into dynamic networks of curved filament bundles^18^ (Fig. 1B, top row; Movie S1, left). Next, we performed this experiment in the presence of defined concentrations of ZapA. At a high concentration of ZapA (6 μM), FtsZ filaments organized into a filament architecture of apparently more aligned and straightened filament bundles (Fig. 1B, lower row; Movie S1, right). This effect became more pronounced over time until the system assembled into a stable steady-state after about 15 min, with an architecture that strongly differed from the pattern found in the absence of ZapA (Fig. 1B). Following these initial observations, we then used automated image analysis methods to quantify the influence of ZapA on the organization of membrane-bound FtsZ filaments.

**Figure 1.**
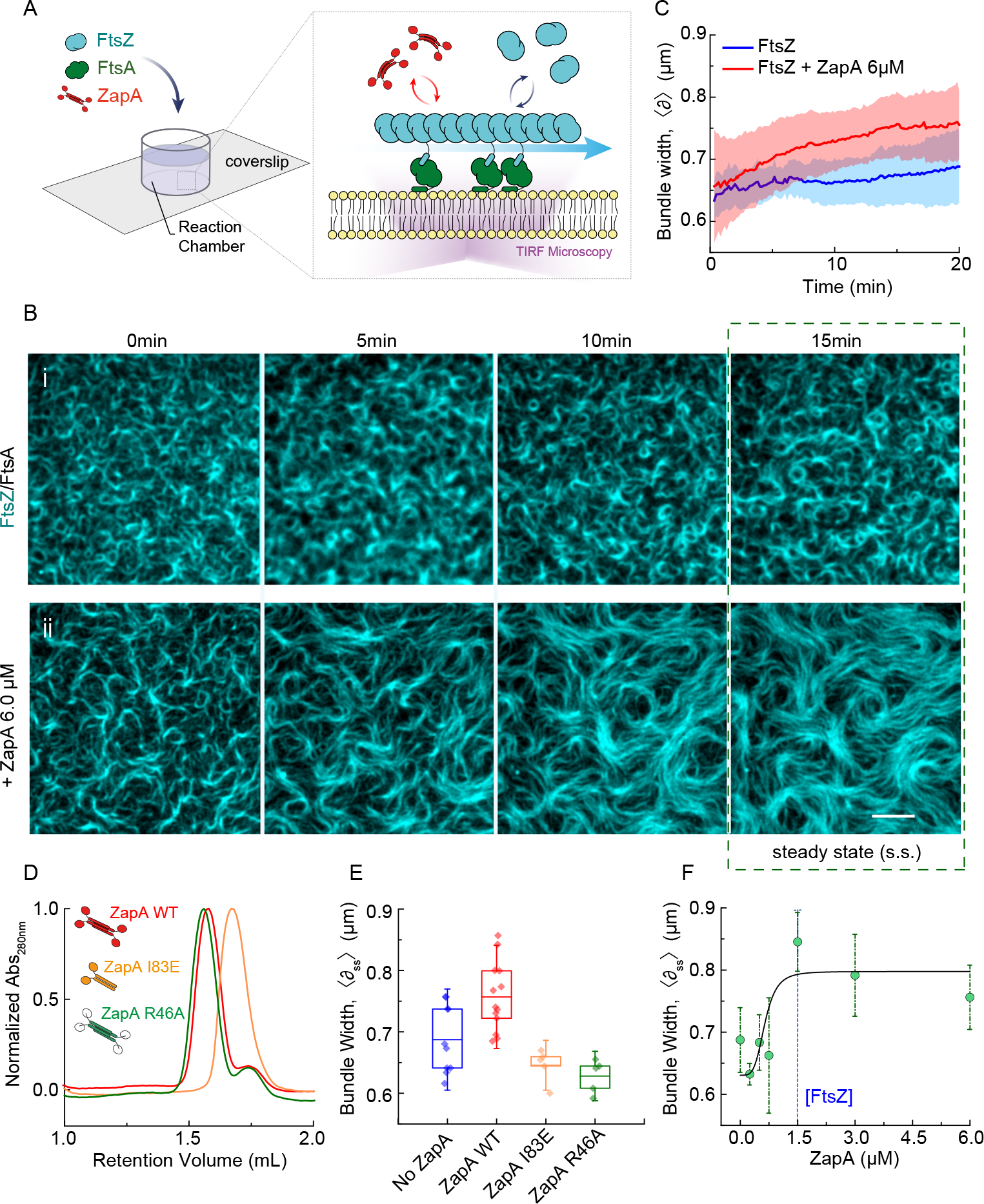
ZapA changes the architecture of membrane-bound FtsZ/FtsA filament networks. **(A)** Illustration of the experimental assay based on a supported lipid bilayer, purified proteins and TIRF microscopy **(B)** Snapshots of FtsZ pattern emerging from its interaction with FtsA alone **(i)** and in the presence of 6μM ZapA **(ii)**. FtsZ (30% Cy5-labelled) and FtsA were kept constant for all experiments (1.5μM and 0.5μM, respectively). Scale bars, 5μm. **(C)** Estimated mean bundle width, 〈*δ*〉, over time for FtsZ/FtsA filament pattern (blue) and with 6μM ZapA (red). Curves depict the mean and standard deviation (shade error bands) of independent experiments (FtsZ, n=13, ZapA, n=14) **(D)** Size-exclusion chromatography results for ZapA wild type (WT) and mutants (ZapA I83E, ZapA R46A) **(E)** Mean bundle width 〈*δ*_*SS*_〉, of membrane-bound FtsZ filaments is increased in the presence of of 6μM ZapA (red) (pval = 0.006) while it was unchanged in the presence of ZapA I83E (yellow) and ZapA R46A (green) (pval > 0.05). A two-tailed Student’s t-test was used to compare the mean values in each condition. **(F)** Change in 〈*δ*_*SS*_〉 with varying concentrations of ZapA. Data shown corresponds to the average bundle width measured between t = 15min and t = 20 min (mean ± s.d). Black line corresponds to Hill equation with n_H_ = 4.12 ± 6.84 and EC_50_ = 0.69 ± 0.31 μM.

As ZapA was previously found to promote lateral interactions between FtsZ protofilaments and the formation of filament bundles when incubated together in solution ^13,16,17^, we first quantified the apparent width of filament structures on the membrane. For this aim, we used an image segmentation approach based on the fluorescence intensity followed by distance mapping (Fig. S1A,B; methods for details). For bundles of membrane-bound FtsZ filaments in the absence of ZapA, our analysis found a mean width of *δ*_*t*=0_ = 0.63 ± 0.33 μm, which stayed roughly constant over time (*δ*_*SS*_ = 0.69 ± 0.06 μm, n = 13; Fig. 1C, blue). In contrast, in the presence of 6.0 μM ZapA, the FtsZ bundle width continuously increased until it plateaued at a slightly higher value of *δ*_*SS*_ = 0.76 ± 0.05 μm (n = 14, Fig. 1C, red) at steady state (t > 15 min). As a control, we prepared ZapA R46A, a mutant of ZapA with an inactive FtsZ binding site (Fig. 1D). As expected, adding this mutant up to a concentration of 6 μM did not change the FtsZ filament pattern and bundles had the same average width as in the absence of ZapA (Fig. 1E, *p* = 0.081; Fig. S1C,D). To confirm that only tetrameric ZapA with two pairs of FtsZ binding sites has this effect on FtsZ filaments, we prepared ZapA I83E, a mutant where the dimer-dimer interaction is disrupted^19^ (Fig. 1D). Even though the presence of ZapA I83E slightly changed the appearance of FtsZ filaments (Fig. S1C, top row), we did not observe a significant degree of bundling as found for the wildtype protein (Fig. 1E; *p* = 9.6E-06; Table S1). Accordingly, and in agreement with previous observations^19^, tetramerization of ZapA is essential for its function.

Next, we were interested in how varying concentrations of wild-type ZapA affects the FtsZ filament pattern (Fig. 1F and Fig. S1E,F). Up to a concentration of ZapA = 0.75 μM, the observed steady-state bundle width was below 0.7 μm and did not change significantly. However, we observed a rapid shift to structures thicker than 0.75 μm at ZapA concentrations higher than 1.5 μM. This value peaked at ZapA = 1.5 μM before it slightly decreased, suggesting a saturation of the system. By fitting a Hill equation to the corresponding data, we found that the behavior of ZapA was highly cooperative with a Hill coefficient of n_H_ = 4.12 ± 6.84 and a half maximal effective concentration of EC_50_ = 0.69 ± 0.31 μM. To summarize, we found that ZapA is able to increase the apparent width of FtsZ filament bundles on the membrane in a cooperative manner.

### ZapA changes the large-scale architecture of FtsZ filament networks

Bundling of FtsZ filaments does not fully characterize the influence of ZapA, as we also observed a dramatic shift in the overall architecture of the filament network. For example, without ZapA, the filament bundles were highly curved, often forming ring-like structures on the membrane. In contrast, the presence of ZapA gave rise to a pattern of straighter filament bundles, where ring-like structures were missing. To better characterize the biochemical activity of ZapA and its influence on FtsZ filament organization, we established a method to obtain a mathematical representation of the filament network (Fig. 2A). Our approach is based on the computation of structure tensors, where each pixel of the image is replaced by a unitary gradient vector describing the average orientation of the local gray scale (methods for details). By applying this method to our time-lapse movies, we obtained an orientation field for every time point of our experiments. The orientation field could then be further analyzed to calculate two independent parameters: the curvature, *κ* (μm^−1^) (Fig. 2B-D) and mean correlation length, *ρ* (μm) (Fig. 2E-F) of the system.

**Figure 2.**
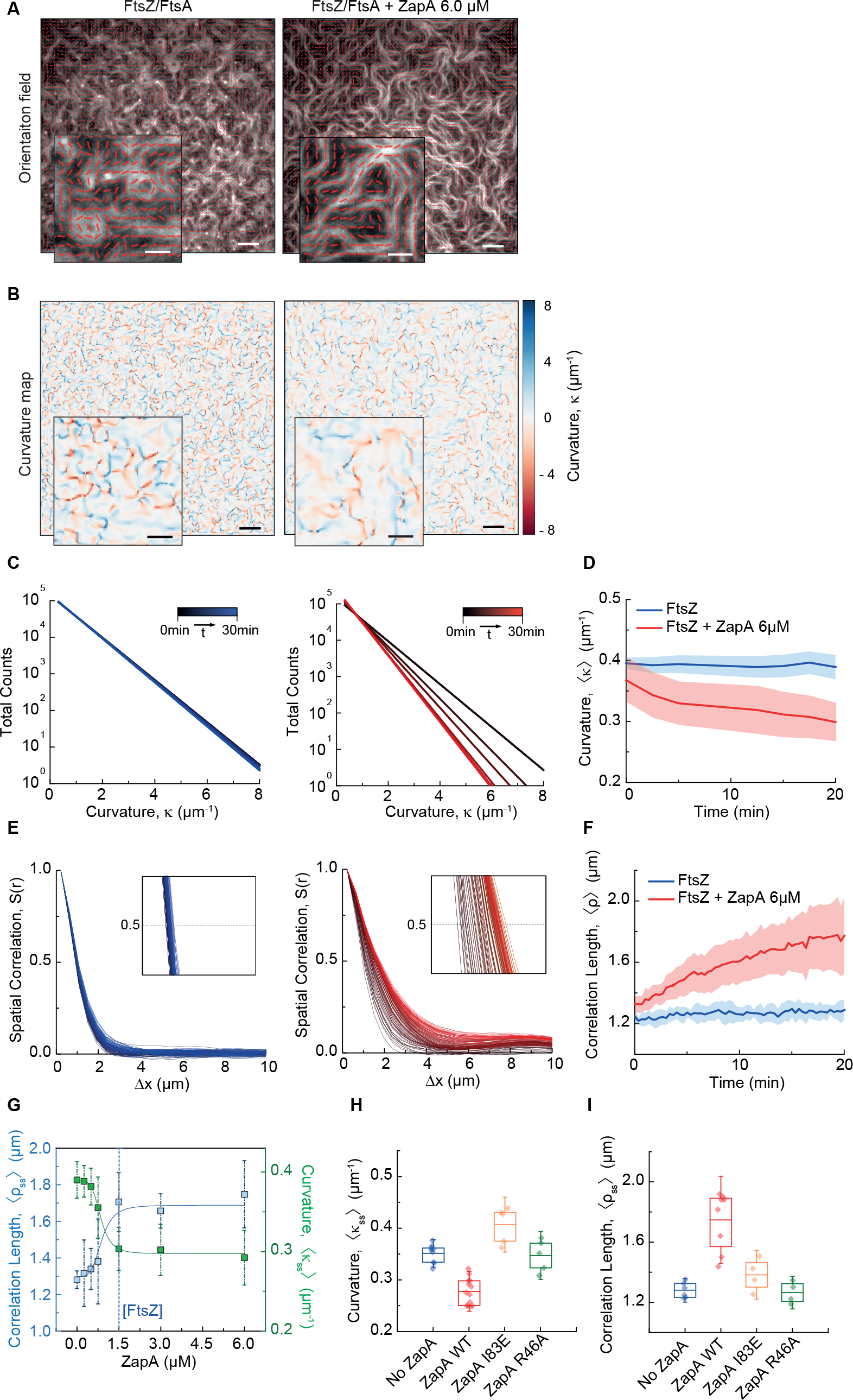
ZapA increases spatial order of FtsZ/FtsA filaments network while decreases its curvature. **(A)** Steady state orientation fields for FtsZ filament network without (left) or with 6.0 μM ZapA (right). **(B)** Curvature maps of orientation fields; white areas correspond to straight filament bundles, red and blue indicate clockwise or counterclockwise rotation, respectively. The two-dimensional distribution of absolute curvature values in each frame was plotted over time. Scale bars: large image, 5 μm; small image, 2μm. **(C)** Time-dependent decrease of FtsZ/FtsA filaments alone (left, black to blue gradient) and with 6μM ZapA (right, black to red gradient) **(D)** The mean curvature, 〈*k*〉, at each time point, was described by the half-time of a mono-exponential decay fit to the curvature distribution values. Curves depict the mean and standard deviation (shade error bands) of independent experiments (FtsZ, n=9, ZapA, n=14). (**E)** Typical spatial correlation function, S(*r*), for an experiment of FtsZ/FtsA network pattern (left, black to blue gradient) or with 6μM ZapA (right, black to red). FtsZ/FtsA filament bundles show overlapping curves in time (black, t=0min, to blue, t=30min) while ZapA promoted a shift to higher correlation lengths within the same time frame (black, t=0min, to red, t=30min), suggesting an increasing spatial order in the system (black, t=0min, to red, t=30min). **(F)** Temporal changes in the correlation length, 〈*ρ*〉, of FtsZ/FtsA filament bundles. Correlation length, given by the intersection line where S(*r*) = 0.5 (insets in **E**) shows that 〈*ρ*〉 remained constant for FtsZ/FtsA filaments alone (blue), but showed a continuous increase the presence of 6μM ZapA. Curves show the mean and standard deviation (shade error bands) of independent experiments (FtsZ, n=6, ZapA, n=8). **(G)** 〈*k*〉 and 〈*ρ*〉 with varying concentrations of ZapA. As for 〈*ρ*〉, ZapA induced a switch-like behavior for 〈*k*_*SS*_〉 and 〈*ρ*_*SS*_〉 (n_H_ = 4.63 ± 0.84 and k_H_ = 0.84 ± 0.05 μM; n_H_ = 4.68 ± 2.84, k_H_ = 0.84 ± 0.17, respectively) with a saturation point at equimolar concentrations with FtsZ. Curves depict the mean and standard deviation (shade error bands) of independent experiments (FtsZ, n=6, ZapA, n=8). **(H,I)** ZapA I83E (yellow) and R46A (green) did not change significantly any of these parameter once again.

From the orientation field in each frame, we obtained a heatmap describing the magnitude of the local curvature at each x and y coordinate (Fig. 2B and Fig. S2A). The mean absolute curvature of the filament network at each time point, *κ*(*t*), was then given by fitting an exponential decay to the curvature distribution (Fig. 2C and Fig. S2A). In the absence of ZapA, FtsZ filaments showed a constant mean absolute curvature of *κ*_*SS*_ = 0.39 ± 0.02 μm^−1^ (Fig. 2C, blue, n = 9). In contrast, in the presence of 6 μM ZapA, *κ*(*t*) decreased over time by about 20% with a plateau at *κ*_*SS*_ = 0.29 ± 0.03 μm^−1^ (Fig. 2C, red, n=13), confirming that the presence of ZapA results in straighter bundles of FtsZ on the membrane.

Next, we used the orientation field to calculate the spatial order of the filament network, which is a measure for how the orientation of the filaments co-vary with one another across the observed membrane area. Specifically, we used the correlation function *S*(*r*) to calculate the distance range *r* over which the orientation angles in each frame share a common direction (methods for details). At a given time, *S*(*r*) showed a decreasing correlation of alignment with increasing distance (Fig. 2E). In the absence of ZapA, the correlation curves obtained for different time points overlapped with each other (Fig. 2E, left; black to blue gradient), showing that spatial order stayed constant during the experiment. The presence of ZapA increases the correlation length, an effect that becomes more pronounced with time, indicating that ZapA increases the spatial order of the system (Fig. 2E, right; black to red gradient). We quantified this time-dependent increase in spatial order by comparing the characteristic correlation length, *ρ*(*t*), which corresponds to the distance at which *S*(*r*) = 0.5 (Fig. 2E insets and Fig. 2F). We found that the presence of ZapA led to a continuous increase in spatial order from *ρ*_*t*_ = 0.97 ± 0.06 μm to *ρ*_*ss*_ = 1.74 ± 0.21 μm at steady state (Fig. 2F, red, n = 8), while *ρ* stayed constant in its absence (Fig. 2F, blue, n = 6).

Next, we measured how increasing concentrations of ZapA affects the curvature and correlation length at steady state, *κ*_SS_ and *ρ*_SS_ (Fig. 2G, Fig. S2B,C). Similar as for the bundle width (Fig. 1F), we found a switch-like transition between two states with Hill coefficients of n_H_ = 4.63 ± 0.84 and n_H_ = 4.68 ± 2.84 and with EC_50_ = 0.84 ± 0.05 and EC_50_ = 0.84 ± 0.17 for *κ*_*ss*_ and *ρ*_*ss*_, respectively. Furthermore, neither ZapA R46A nor I83E had the same effect on the FtsZ filament network (Fig. 2H,I; Table S1). These results show that the system exists in two states: a state of high curvature and low spatial order, at ZapA concentrations below 0.8 μM, and a state of high spatial order and low curvature at higher concentrations.

### ZapA increases persistence of the membrane-bound FtsZ filament network

Z-rings in cells lacking ZapA not only show loose arrangement, but are also highly dynamic, transitioning back and forth between multiple locations in the cell^20^. In contrast, the Z-ring in wildtype cells usually persists at the same position during the cell cycle. Consistent with these observations, we realized that FtsZ filament bundles continuously reorganized in the absence of ZapA, but appeared much more static when ZapA was present. To quantify the degree of reorganization, we performed temporal autocorrelation analysis. This method measures the similarity of fluorescence images as a function of a time lag Δt between different frames (Fig. 3A). We found that the corresponding autocorrelation function, θ(Δt), rapidly decayed in the absence of ZapA, consistent with a rapidly rearranging pattern (Fig. 3B, blue curve). In the presence of 6μM of ZapA, however, this decay was about 4-fold slower (Fig. 3B, red curve). We defined the characteristic correlation time, *τ* for varying concentrations of ZapA by the halftime of a mono-exponential fit to the correlation curves. Similar to bundle width, curvature and correlation length, *τ* displayed a two-state behavior: below a critical concentration of ZapA the pattern showed fast reorganization and consequently short correlation times, but the system switched to slow reorganization and long correlation times at saturating ZapA concentrations with a Hill coefficient of 4.89 ± 5.13 and EC_50_ = 0.79 ± 0.21 μM (Fig. 3C). ZapA R46A, which does not bind to FtsZ, did not affect reorganization dynamics (Fig. 3D; *p* = 0.42), while ZapA I83E increased slightly, but not-significantly the correlation time (Fig. 3D; *p* = 0.08). This shows that tetrameric ZapA not only affects the architecture of the membrane-bound filament network, but also its reorganization dynamics in a highly cooperative manner.

**Figure 3.**
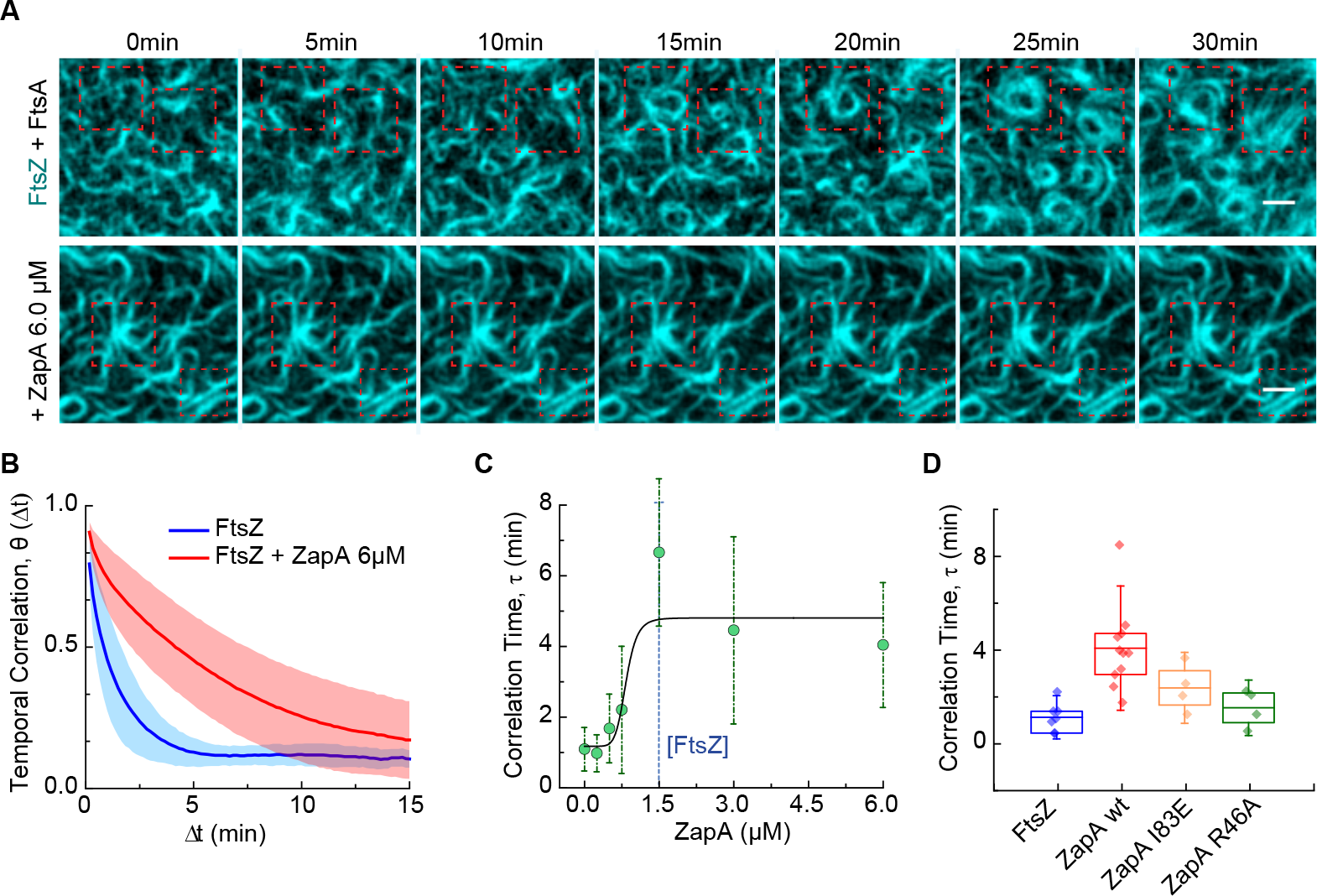
ZapA increases persistence of FtsZ filaments. **(A)** In the absence of ZapA (upper panel), the FtsZ filament pattern constantly reorganized and ring-like structures formed only transiently (red dashed squares). In contrast, in the presence of ZapA (lower panel) FtsZ assemblies were much more persistent. Scale bars, 2 μm **(B)** The temporal autocorrelation function of the FtsZ filament network decayed quickly without ZapA (blue), but slowly in the presence of 6.0 μM (red). Each curve represents the mean ± s.d (error bands) of independent experiments (FtsZ, n=7, ZapA, n=11). **(C)** The characteristic correlation time, *τ*, as a function of ZapA concentration. The autocorrelation curves were fitted to a mono-exponential decay and *τ* was given by the half-time of each decay. Data shown corresponds do the average / from individual experiments (mean ± standard deviation). **(D)** ZapA I83E (yellow) and R46A (green) did not change the persistence of FtsZ filaments.

Together, our *in vitro* experiments and quantitative image analysis show that ZapA promotes an ultrasensitive switch in filament architecture and reorganization dynamics. In all instances, this switch happens at ZapA concentrations around 0.75 μM, before the system saturates at ZapA concentrations similar to FtsZ. Interestingly, this correlation is consistent with ZapA tetramers having four FtsZ binding sites.

### ZapA does not change FtsZ polymerization dynamics

So far, our analysis revealed that ZapA strongly increases the spatial order of the membrane-bound FtsZ filament network, and slows down reorganization dynamics. This apparent increase in filament stability, however, could perturb the spatiotemporal dynamics of cell wall synthases *in vivo*, as their motion is driven by treadmilling. Knocking out individual Zap proteins *in vivo* did not show any significant change in the FtsZ treadmilling velocity^5^. However, due to their overlapping functions, it is difficult to rule out a possible compensation by other FtsZ associated proteins in these mutants. Accordingly, we wondered if FtsZ crosslinking by ZapA influences the polymerization dynamics of membrane-bound filaments. To quantify the treadmilling velocity in the absence and presence of ZapA, we first constructed differential time-lapse movies, where we calculated the intensity differences between frames separated by a constant time delay (Fig. 4A and Fig. S3; Movie S2). This step yielded new time-lapse movies of moving speckles that represent either the growth or shrinkage of filament bundles in a given time (methods for details).

**Figure 4.**
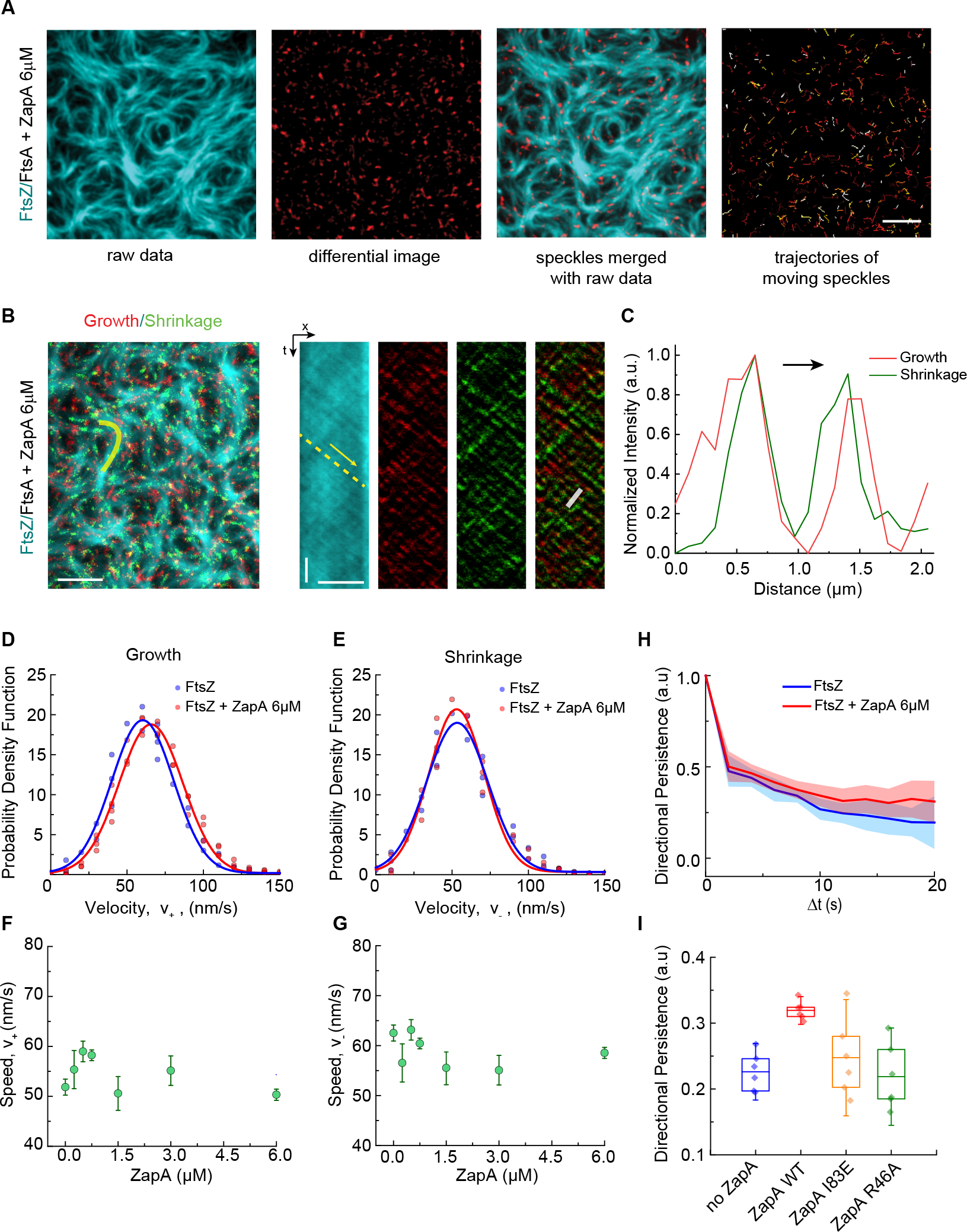
ZapA has no effect on FtsZ treadmilling dynamics. **(A)** Differential TIRF images allows for tracking polymerization (growth) and depolymerization (shrinkage) of FtsZ treadmilling. **(B)** Raw image of FtsZ filament network in the presence of 6 μM ZapA overlaid with the respective differential image showing growing (red) and shrinking (green) ends. Kymographs on the right show position fluctuations in time along the yellow line. Scale bars, t = 5 s, × = 5 μm. **(C)** Plot profile along the white line in the Kymograph represents an average of intensities of growing and shrinking ends from multiple time points along the treadmilling path. Common direction of growth and shrinking reveals the parallel organization of filament bundles, colocalization indicates the presence of short protofilaments. **(D)** Velocities distribution of filament growth (FtsZ/FtsA, ***ν* +** = 62.5 ± 4.5 nm/s, FtsZ/FtsA + ZapA, ***ν* +** = 59.9 ± 3.5 nm/s) and **(E)** shrinkage (FtsZ/FtsA, ***ν*−** = 51.8 ± 6.0 nm/s, FtsZ/FtsA + ZapA, ***ν*−** = 50.3 ± 3.3 nm/s). Both show a normal distribution without any significant difference in the mean. (**F, G)** Increasing concentrations of ZapA did not affect the mean velocity of FtsZ filaments at both ends (n = 45). **(H)** Velocity autocorrelation revealed that directionality was significantly increased (pval < 0.001) in the presence of 6μM ZapA (***ν***_corr_(FtsZ) = 0.226 ± 0.029 and ***ν***_corr_ (ZapA) = 0.319 ± 0.014 for Δt = 10 - 20s). **(I)** ZapA mutants did not change treadmilling directionality (pval > 0.05).

First, as ZapA was suggested to promote an antiparallel alignment of FtsZ filaments, we were interested if the presence of ZapA leads to a change of filament orientation^12^. Comparing the speckles between experiments with and without ZapA, we did not observe any significant difference, neither in shape nor intensity (Fig. S3A). This shows that the orientation of filaments in ZapA-induced bundles was similar as in bundles formed due to intrinsic lateral interactions between FtsZ filaments. Furthermore, we observed that fluorescent speckles corresponding to growth and shrinkage along the bundle roughly co-localized, and moved in a common direction with similar velocity (Fig. 4B,C). This observation is consistent with a polar orientation of FtsZ filaments on the membrane and indicates that the mean filament length is shorter than the resolution of our fluorescence microscope (Fig. S3B).

Importantly, as these speckles can be automatically detected by particle tracking methods, differential imaging allows us to quantify the growth (***ν*_+_**) and shrinkage velocities (***ν*_−_**), of thousands of treadmilling trajectories simultaneously (methods for details). Using this approach, we found ***ν*_+_** and ***ν*_−_** to be normally distributed with similar mean values consistent with treadmilling behavior (***ν*_+_** = 62.5 ± 4.5 nm/s, n = 4232 tracks, 10 samples; ***ν*_−_** = 51.84 ± 5.9 nm/s, n = 6302 tracks, 10 samples) (Fig. 4D,E, blue). These values were unaffected in the presence of excess ZapA (6.0 μM, ***ν*_+_** = 58.8 ± 3.3 nm/s and ***ν*_−_** = 54.3 ± 3.5 nm/s) (Fig. 4D,E, red) and at all other ZapA concentrations tested in our experiments (Fig. 4F,G).

The velocity autocorrelation along each treadmilling trajectory provides information about the local directional persistence of treadmilling (Fig.4H; methods for details). At short times the velocity correlation in the absence of ZapA is always positive, consistent with persistent treadmilling filaments. However, we found that in the presence of excess concentrations of ZapA (Fig.4H, red) the velocity remained correlated for a longer time, consistent with straighter treadmilling tracks (Fig.4H, blue). Again, neither 6.0 μM ZapA I83E nor ZapA R46A showed an effect on the directional persistence of FtsZ treadmilling (Fig. 4I; *p* = 0.45 and *p* = 0.76, respectively; Table S1).

Together, these experiments and analyses show that despite its strong effect on the architecture and reorganization dynamics of the FtsZ filament network, ZapA does not slow down or enhance the underlying polymerization dynamics.

### ZapA binds FtsZ only transiently

Next, we sought to better understand how ZapA could change the architecture of FtsZ filaments without slowing down their treadmilling dynamics. To this end, we prepared fluorescently labeled ZapA and imaged its behavior simultaneously with FtsZ. We found that ZapA co-localized with FtsZ bundles on the membrane, however with a more discontinuous appearance (Fig. 5A; Movie S3) suggesting highly dynamic, transient binding of ZapA to the filaments.

**Figure 5.**
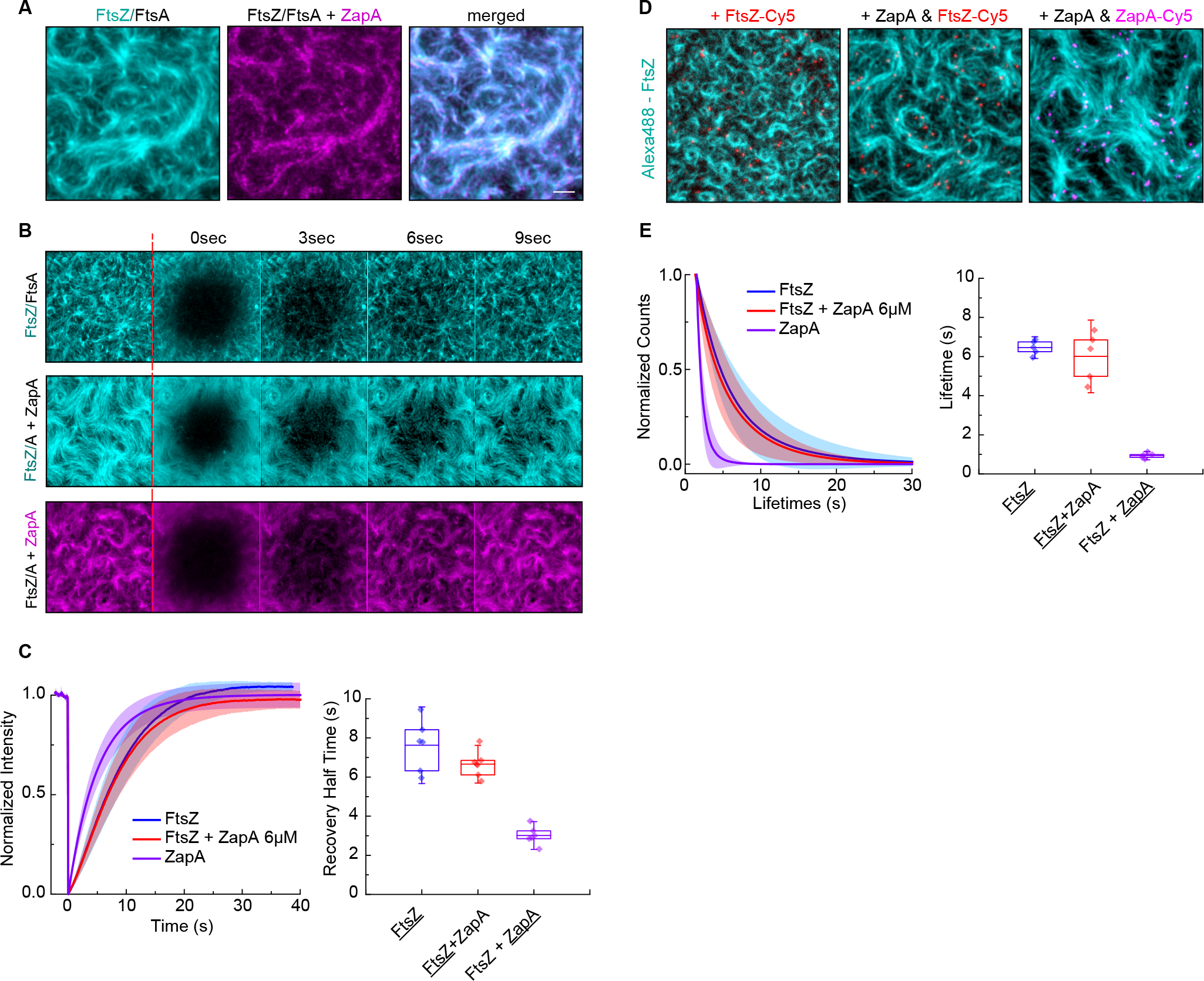
ZapA has a faster turnover rate than FtsZ. **(A)** Dual-color TIRF micrograph showing co-localization of Alexa488-FtsZ (cyan) and ZapA-Cy5 (magenta). Single particle tracking (SPT) of FtsZ (red) and ZapA (magenta). **(B)** Scale bars, 5 μm **(C)** Lifetime distribution of FtsZ does not change significantly with ZapA (FtsZ-Cy5 = 6.59 ± 0.28 s, blue, n = 5; FtsZ-Cy5/FtsA + ZapA = 6.01 ± 1.24 s, red, n = 6). In contrast, the lifetime of ZapA was much shorter (ZapA-Cy5 = 0.95 ± 0.14, n = 5). **(D)** Snapshots showing fluorescence recover after photobleaching (FRAP) experiments. **(E)** Half-time recovery of FtsZ was not affected by 6μM ZapA (7.63 ± 1.30 s and 6.66 ± 0.64 s, respectively), while Cy5-labelled ZapA showed a much faster recovery (ZapA-Cy5 = 3.01 ± 0.47, n = 6).

We then analyzed the turnover rate of FtsZ and ZapA using fluorescence recovery after photobleaching (FRAP) experiments (Fig. 5B; Movie S4). For FtsZ, we found a mean recovery half-time of 7.63 ± 1.30 s (n = 6) in the absence of ZapA and 6.66 ± 0.64 s (n = 7) at 6.0 μM ZapA, consistent with our observation that treadmilling was unchanged (Fig. 5C; *p* = 0.47). In contrast, ZapA itself showed a faster turnover with a recovery half-time of only 3.01 ± 0.47 s (n = 6). This difference in turnover is similar to the one found in living cells^8^.

To further corroborate these results, we performed single molecule experiments, where we added small amounts of a Cy5-labelled protein, either ZapA or FtsZ, to a background of Alexa488-labelled FtsZ (Fig. 5D; Movie S5). This not only allowed us to analyze the lifetime of FtsZ monomers in the treadmilling filaments, but also the recruitment of ZapA from solution. In agreement with our FRAP experiments, we found no difference in the lifetimes of FtsZ with and without ZapA (FtsZ, 6.59 ± 0.28 s, n = 5; FtsZ + ZapA, n = 6.01 ± 1.24 s, n=6; *p* = 0.14) (Fig. 5E, blue and red, respectively). Again, ZapA showed a faster turnover compared to FtsZ (0.94 ± 0.14 s, n = 5; Fig. 5D, purple; *p* = 1.44E-04; Table S1). Together with the treadmilling velocity, we can now provide an estimate of the mean length of filaments, which is given as the product of lifetime with the treadmilling velocity. We found an FtsZ filament length between 341.6 ± 50.0 and 412.0 ± 37.7 nm, when only FtsA was present, and between 302.4 ± 72.9 and 351.8 ± 76.2 nm in the presence of ZapA, showing that ZapA does not change the filament length. These values obtained in our *in vitro* experiments are about three times longer than what has been suggested for filaments *in vivo*^21^.

These results show that despite its strong influence on their organization, ZapA binds only transiently to FtsZ filaments. We believe that the difference in residence times of FtsZ and ZapA can explain how ZapA can increase the persistence and order of FtsZ filaments without compromising the treadmilling dynamics.

## Discussion

FtsZ has long been known as the main organizer of bacterial cell division. Recent studies have shown that FtsZ plays two roles: First, its polymerization into the Z-ring defines the location of division in the cell. Second, FtsZ treadmilling was found to drive a circumferential motion of peptidoglycan synthases, which is required for the homogeneous distribution of cell wall synthesis at the division site. Given the importance of FtsZ polymerization dynamics for cytokinesis, the question arises how the robustness and precision of cell division is achieved, while FtsZ filaments are continuously turning over. FtsZ-associated proteins were known to contribute to the stability of the Z-ring and consequently that of the cell division machinery, the underlying mechanism however was not clear.

In this study, we show that ZapA aligns membrane-bound FtsZ filaments in a polar, parallel orientation. This results in a straightening and stabilization of filament bundles, and consequently an increase in the spatial order and spatiotemporal persistence of the filament network. Importantly, ZapA did not change the treadmilling velocity of FtsZ filaments (see Table 1 for a summary of results). Together, these observations lead us to conclude that ZapA is able to increase the precision and robustness of the Z-ring, without compromising the function of the cell division machinery (Fig. 6).

**Figure 6.**
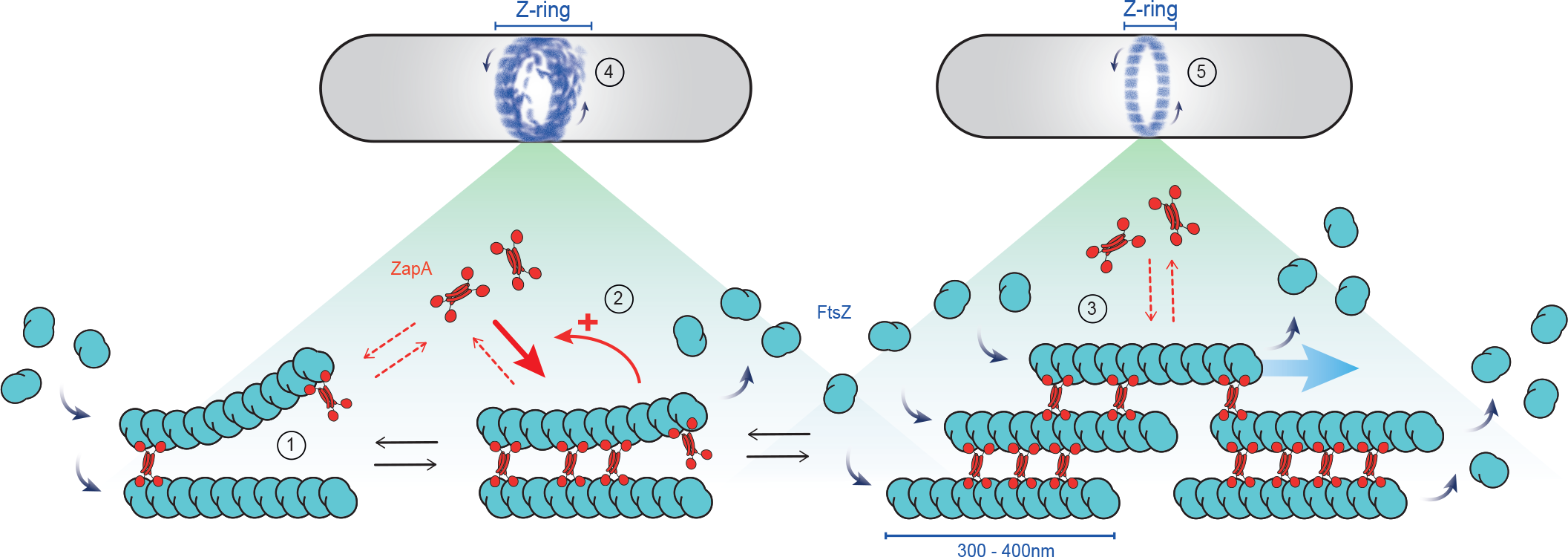
Cooperative binding of ZapA on FtsZ/FtsA filaments. Binding of ZapA aligns treadmilling FtsZ filaments **(1)**, which in turn facilitates its binding, giving rise to cooperativity **(2).** At saturation, four molecules of FtsZ bind to one tetramer of ZapA (1:1 monomer ratio, [FtsZ] = [ZapA]) in a parallel fashion **(3)** increasing the bundle width, persistence and alignment of FtsZ filaments. Treadmilling is not affected due to the transient binding of ZapA **(4).** This process leads to a shift from a network of less ordered and highly dynamic FtsZ filaments into a more well-defined track, acting as an additional fine-tune mechanism for a proper Z-ring alignment without compromising treadmilling dynamics.

**Table 1.** Summary of all the parameters retrieved from the analyses. Weighted mean and respective standard deviation of each parameter for all ZapA concentrations combined, either below 0.75 μM (low ZapA) or above 1.5 μM (high ZapA). This shows how much the system changed between the two states.

|  |  | Low ZapA ( < 0.5uM) |  |  | High ZapA ( > 1.5uM) |  |  |  |  |
| --- | --- | --- | --- | --- | --- | --- | --- | --- | --- |
| Parameter | Observation | mean | s.d | n | mean | s.d | n | % Change | Figure |
| Bundle width | ↑ | 0.68 | 0.02 | 18 | 0.78 | 0.02 | 40 | 15% | Fig.1 |
| Spatial Order | ↑ | 1.31 | 0.30 | 14 | 1.71 | 0.04 | 21 | 30% | Fig.2 |
| Curvature | ↓ | 0.39 | 0.00 | 17 | 0.30 | 0.01 | 34 | 23% | Fig.2 |
| Persistence | ↑ | 1.26 | 0.30 | 15 | 5.23 | 1.17 | 34 | 314% | Fig.3 |
| Growth Rate | = | 62.2 | 2.5 | 18 | 57.0 | 1.61 | 23 | n.s | Fig.4 |
| Shrinkage Rate | = | 55.6 | 0.9 | 18 | 53.4 | 1.43 | 23 | n.s | Fig.4 |
| Directionality | ↑ | 0.23 | 0.03 | 10 | 0.32 | 0.01 | 12 | 41% | Fig.4 |
| Lifetime | = | 6.46 | 0.37 | 5 | 6.01 | 0.01 | 5 | n.s | Fig.5 |
| FRAP Half-Time | = | 7.63 | 1.30 | 6 | 6.66 | 0.64 | 7 | n.s | Fig.5 |

| Hill Equation Fitting | EC50 | Hill Coefficient | R<sup>2</sup> |
| --- | --- | --- | --- |
| Bundle Width | 0.64 ± 0.28 | 4.12 ± 6.84 | 0.68531 |
| Correlation length | 0.84 ± 0.17 | 4.68 ± 2.84 | 0.95684 |
| Curvature | 0.85 ± 0.17 | 4.63 ± 0.84 | 0.99581 |
| Correlation Time | 0.79 ± 0.21 | 4.89 ± 5.13 | 0.81486 |

Furthermore, we found that ZapA promotes a switch-like transition between a state of low order and fast reorganization dynamics to a state of high order and slow reorganization. What could be the reason for the observed cooperativity? Due to its central four-helix bundle, the ZapA tetramer can be assumed to have a rigid structure, which promotes the formation of straight filament bundles as observed in previous studies^13^. Furthermore, we expect the ZapA tetramer to bind with a higher affinity to filaments aligned in parallel, where all binding sites are in contact with FtsZ. Accordingly, binding to and aligning filaments can increase the affinity of ZapA towards FtsZ and therefore stimulate its own recruitment (Fig. 6).

Our experiments show that FtsZ, FtsA and ZapA self-organize into a highly ordered cytoskeletal structure. How does this system compare to filament networks of other cytoskeletal proteins? Similar to gels of actin filaments and molecular motors, the structures described here exist out of equilibrium. In contrast to these active gels, however, the FtsZ filament network is driven out of equilibrium due to the continuous consumption of energy during treadmilling and not by the activity of motor proteins. Furthermore, the function of actin bundling proteins is to modulate the architecture and mechanical properties of actin filament networks, which can also affect actin polymerization dynamics^22^. In contrast, polymerization and depolymerization rates of FtsZ stay constant even at saturating concentrations of ZapA. Accordingly, we believe that the cytoskeletal networks of FtsZ, FtsA and ZapA represent a novel and distinct form of active biological material despite the apparent absence of mechanical stresses that are usually present in gels composed of filaments and molecular motors^23-25^.

## Acknowledgements

We thank all Loose lab members for support and useful discussions, to the life sciences facility (LSF) and the bioimaging facility (BIF) of IST Austria for assistance with general equipment and TIRF microscopy, respectively, and to Georgia Squyres (Harvard University) for critical revision of results and manuscript before publication; This work was supported by a European Research Council (ERC) grant awarded to Martin Loose (ERC-2015-StG-679239) and a Boehringer Ingelheim Fonds (BIF) PhD fellowship awarded to Paulo Caldas.

## Author contributions

**Project Management**: M.L; **Conceptualization & Design**: M.L and P.C; **Experimental Data Collection**: P.C and M.L; **Curvature Analysis**: N.B and P.C.; **Spatial Order Analysis**: D.P, M.L and P.C; **Treadmilling speed and further image analysis**: M.L, P.C and J.B; **Data interpretation**: M.L, J.B and P.C; **Data Visualization**: P.C; **Manuscript Drafting**: M.L and P.C; **Critical Revision of the Manuscript**: All authors; Final version was read by all authors and approved to be published.

## Competing interests

The authors declare no competing interests.

## Material and Methods

### Reagents and Chemicals

Phospholipids used in this paper, DOPC (1,2-dioleoyl-sn-glycero-3-phosphocholine) and DOPG (1,2-dioleoyl-snglycero-3-phospho-(1′-racglycerol)), were purchased from Avanti Polar Lipids (Alabaster, AL) and kept at −20°C as 25mg/ml stock solutions in chloroform; Sulfo-Cyanine-5-maleimide (Cy5) was acquired from Lumiprobe and Alexa Fluor^®^ 488 C5-Maleimide (Alexa488) was acquired from ThermoFisher Scientific; Nucleotides were acquired from ThermoFisher Scientific or Jena Bioscience; Precision cover glasses for the homebuilt chambers were obtained from VWR (thickness No. 1.5H, 24 × 50); *E.coli* strains were obtained from Lucigen; Strep-Tactin resin was acquired from Iba lifesciences and Nickel resins were purchased from ThermoFisher Scientific (HisPur™ Ni-NTA resin) or Macherey-Nagel (Protino Ni-TED resin); All the remaining reactants and salts were obtained from Sigma, Merck, or Invitrogen and were of analytic or spectroscopic grade.

### Protein Biochemistry

#### Purification and labelling of FtsZ

FtsZ (UP P0A9A6) was cloned into a pTB146-derived vector which attached a N-terminal His_6_-SUMO tag plus seven additional amino acids (AEGCGEL) that provide a cysteine residue for further fluorescent labelling (pML45, His_6_-SUMO-GCG-FtsZ). *E.coli* C41 (DE3) cells were transformed with pML45 and grown in TB medium supplemented with ampicillin at 37°C, until cells reached an OD600 of 0.8. After the addition of IPTG to a final concentration of 1mM, cells grew for 5h at 37°C. Cells were harvested by centrifugation, pellets were frozen in liquid nitrogen and kept at −80°C until further use.

For purification, pellets were thawed and resuspended in FtsZ buffer (50 mM Tris-HCl pH 8.0, 500mM KCl, 2mM β-mercaptoethanol, 10% glycerol) supplemented with 20mM imidazole and cOmplete EDTA-free protease inhibitors cocktail (1tablet/50ml, Roche)) followed by incubation at 4°C for 15 min. Cells were lysed using a cell disruptor (Constant Systems) at a pressure 1.36kbar and incubated with 1mg/ml DNase I (Sigma-Aldrich) and 2.5mM MgCl_2_ for 15min. The lysate was then centrifuged (30 min, 60,000xg, 4°C) and the supernatant was incubated with nickel agarose resin (HisPur Ni-NTA resin, Thermo Scientific) for 60 min at 4°C. The resin was extensively washed with FtsZ buffer supplemented with 20mM imidazole and 30mM imidazole. The fusion protein was eluted with FtsZ buffer supplemented with 250 mM imidazole. Fractions were evaluated by SDS-PAGE (stained with Coomasie Blue) and peak fractions containing His_6_-SUMO-GCG-FtsZ were pooled and incubated with His_6_-tagged SUMO protease (His_6_-Ulp1) during an overnight dialysis into FtsZ cleavage buffer (50mM Tris-HCl pH 8.0, 300mM KCl and 10% glycerol). The digested sample was passed several times through Ni-NTA resin, to remove His_6_ containing molecules. The flow through was collected and active protein was enriched by sedimentation of FtsZ filaments. For this, the protein was dialyzed into FtsZ polymerization buffer (50 mM PIPES pH 6.7, 10 mM MgCl_2_) and incubated with CaCl_2_ and GTP for 15 min at 30°C. The solution was then centrifuged (2 min, 20,000xg, RT) and a clear gel-like pellet containing polymeric FtsZ was obtained. The pellet was resuspended into FtsZ storage buffer (50mM Tris-HCl pH 7.4, 50mM KCl, 1mM EDTA and 10% glycerol) and incubated with 100×molar excess of 7 Tris(2-carboxyethyl)phosphine hydrochloride (TCEP) for 20 min at RT for cysteine-labelling. Five times molar excess of a thiol-reactive dye (Alexa488 or Cy5–maleimide) was added to the solution and incubated overnight at 4°C during dialysis into FtsZ storage buffer. Finally, labelled FtsZ was loaded on a PD10 desalting column to remove CaCl_2_, GTP and free dye. Purified FtsZ was aliquoted, flash frozen in liquid nitrogen and kept at −80°C until usage.

#### Purification of FtsA

The gene coding for FtsA (UP P0ABH0) was cloned into a modified pTB146 vector. The resulting vector, pML60, encodes for FtsA with an N-terminal His_6_-SUMO-pentaglycine tag. *E.coli* C41 (DE3) cells were transformed with pML60 and grown in 2xYT medium supplemented with ampicillin, at 37°C, until they reached an OD600 of 0.6-0.8. After the addition of IPTG to a final concentration of 1mM, cells grew overnight at 18°C. Cells were harvested by centrifugation, pellets were frozen in liquid nitrogen and kept at −80°C until further use.

For purification, cells were thawed and resuspended in FtsA buffer (50 mM Tris-HCl pH 8.0, 500mM KCl, 10mM MgCl_2_) supplemented with 1 mg/ml lysozyme, cOmplete EDTA-free protease inhibitors cocktail (1 tablet/50ml, Roche), 1 mg/ml DNase (Sigma-Aldrich) and 0.5mM DTT. Cells were then lysed using a cell disruptor (Constant Systems) at a pressure of 1.36kbar, centrifuged (60,000xg, 45min at 4°C) and the supernatant was incubated with Nickel agarose beads (Protino Ni-TED, Macherey-Nagel) for 1h at 4°C.

The resin was extensively washed with FtsA buffer and FtsA buffer supplemented with 5mM imidazole. The fusion protein was then eluted with FtsA buffer supplemented with 250mM imidazole and fractions were evaluated by SDS-PAGE (stained with Coomassie Blue). Peak fractions containing His-SUMO-GGGGG-FtsA were pooled and the buffer was exchanged into FtsA storage buffer (50 mM Tris-HCl pH 8.0, 500 mM KCl, 10 mM MgCl_2_, 0.5 mM DTT, 20% glycerol). The His_6_-SUMO tag was cleaved by incubating His-SUMO-GGGGG-FtsA with His_6_-Ulp1 for 90min at 30°C. The sample was then passed several times through Protino Ni-TED resin, previously equilibrated with FtsA storage buffer, to remove His_6_-containing molecules. Protein was aliquoted, flash frozen in liquid nitrogen and kept at −80°C until usage.

#### Purification of ZapA and Mutants

ZapA (UP P0ADS2) was cloned into a modified pTB146 vector which attached an N-terminal Twinstrep-SUMO tag (pML130, Twinstrep-SUMO-ZapA). *E.coli* BL21 (DE3) cells were transformed with pML130 and grown in TB medium supplemented with ampicillin, at 37°C, until cells reached an OD600 of 0.6-0.8. After the addition of IPTG to a final concentration of 1 mM, cells grew for 4h at 37°C. Cultures were harvested, and pellets were frozen in liquid nitrogen and kept at −80°C until further use.

For purification, cells were thawed and incubated with ZapA buffer (50 mM HEPES pH 7.5, 300mM KCl, 10% glycerol) supplemented with 2mM β-mercaptoethanol and cOmplete EDTA-free protease inhibitor tablets (1tablet/50mL, Roche Diagnostics) for 15min at 4°C. Cells were then lysed using a cell disruptor and incubated in the presence of 1mg/ml of DNase (Sigma-Aldrich) and 2.5mM MgCl_2_ for 15min. The lysate was centrifuged (30 min, 60,000xg, 4°C) and the supernatant was incubated with strep-tactin resin (Strep-Tactin^®^ resin, Iba Lifesciences) for 30min at 4°C. Beads were extensively washed with ZapA buffer and the fusion protein was eluted using ZapA buffer supplemented with 50mM biotin. Peak fractions of Twinstrep-SUMO-ZapA were identified by SDS-PAGE (stained with Coomassie Blue) and pooled. The Twinstrep-SUMO tag was cleaved with His_6_-Ulp1 during an overnight dialysis into ZapA storage buffer (50mM Tris-HCl pH 7.5, 50 mM KCl, 10% Glycerol). The solution was then passed through Ni-NTA and strep-tactin resins to remove His_6_-Ulp1 and free twinstrep-SUMO tag, respectively. ZapA was aliquoted, flash frozen in liquid nitrogen and kept at −80°C until usage.

For labeling purposes, ZapA was additionally cloned into a vector which attached 5 residues (LPETG) to the C-terminus of the protein, pMAR11b (TwinStrepSUMO-ZapA-LPETG). ZapA-LPETG was expressed and purified as the wildtype protein followed by sortase-mediated labelling ^26^. Specifically, ZapA-LPETG was incubated with 0.5 mM of labelled peptide GGGC-Cy5 and 10μM Sortase 7M (purified using pET30b-7M SrtA, a gift from Hidde Ploegh, Addgene plasmid #51141) in a final concentration of 50 μM. The reaction was carried during an overnight dialysis into ZapA storage buffer at 4°C. Sample was further purified from free labelled peptide and Sortase by size-exclusion chromatography using a Hi/Load 16/600 Superdex 75 column (GE Healthcare) previously equilibrated with ZapA storage buffer. Labelled ZapA was collected, aliquoted and flash frozen in liquid nitrogen. ZapA mutants, R46A and I83E, were obtained by site-directed mutagenesis (pMAR9 and pMAR10b, respectively) and purified following the same procedure as the wild type protein.

### Self-organization assay and Imaging

#### Preparation of small unilamellar vesicles (SUVs)

DOPC and DOPG dissolved in chloroform were transferred into a glass vial to a final concentration of 5 mM (70:30 molarity ratio). Using a stream of nitrogen, the mixture was dried to form a thin film of lipids and transferred to a vacuum desiccator for 3h to completely remove any residual chloroform. The lipids were then rehydrated in Supported Lipid Bilayer (SLB) buffer (50mM Tris-HCl pH 7.5, 300mM KCl) and incubated at 37°C for 30 min. The lipid film was then completely resuspended by vortexing vigorously to obtain multilamellar vesicles of different sizes and frozen using liquid nitrogen. To obtain small unilamellar vesicles (SUVs), the suspension was first subjected to five freeze-thaw cycles followed by tip sonication. Vesicles were kept at −20°C in small aliquots of 30μl up to two months.

#### Preparation of self-made reaction chamber

Glass coverslips were cleaned by sonicating in 2% Hellmanex II solution (Hellma) for at least 15 min, followed by extensive washing and sonication in milliQ H_2_O before storage in 95% reagent-grade ethanol for up to one week. Before use, glass coverslips were blown dry with compressed air and cleaned in an air plasma for 15 min. The reaction chamber was prepared by attaching a plastic ring (top of Eppendorf tube) on a cleaned glass coverslip using ultraviolet glue (Norland optical adhesive 88).

#### Preparation of supported lipid bilayer (SLB)

To form a supported lipid bilayer, a SUV suspension was diluted to 1mM in SLB buffer (50mM Tris-HCl pH 7.5, 300mM KCl) and CaCl_2_ was added to a final concentration of 2mM. From this mix, 50 μl were added to each self-made reaction chamber. Vesicles adsorb to the surface, rupture and fuse with the clean hydrophilic glass to form a flat bilayer, which is further facilitated by the presence of CaCl_2_. After 1h of incubation at room temperature, 50 μl of SLB buffer was added to each chamber (for a final volume of 100μl) and sample was rinsed several times with 200 μL of reaction buffer (50mM Tris-HCl pH 7.5, 150mM KCl, 5mM MgCl_2_).

For the self-organization assay, all experiments were performed by first adding an oxygen scavenger system to the reaction chamber (0.2% d-Glucose, 0.016 mg/ml glucose oxidase, 0.002mg/ml catalase, 1mM dithiothreitol and 0.25mg/mL trolox) to prevent photobleaching. Then, FtsZ (with 30% Cy5- or Alexa488-labeled protein) and FtsA were added to the reaction chamber in a final concentration of 1.5 μM and 0.5 μM, respectively. To trigger polymerization and membrane recruitment of FtsZ, ATP and GTP were added to the system to a final concentration of 4 mM. ZapA was added in different concentration ranges before the addition of GTP.

#### Total internal reflection fluorescence microscopy (TIRFM)

All experiments were performed on an Inverted multipoint total internal reflection fluorescence (TIRF) microscope (TILL Photonics) equipped with dual camera TIRF objectives (Andor 897straight (X-8449) 512×512 pixel and Andor 897 (X-8533) 512×512 pixel) and an image splitter (Andor Tucam) equipped with a long pass of 580 and 640nm. Alexa488 and Cy5 dyes were excited using 488nm and 642nm laser lines, respectively, and the emitted light was filtered by an Andromeda quad-band bandpass filter. Images were typically obtained every 2s, with 50ms exposure to minimize photobleaching and using a 100x Olympus TIRF (NA = 1.49 DIC) objective.

### Image analysis and processing

For image processing, image stacks were imported using FIJI software ^27^. For image analysis and for visualization (supplemental movies) acquired time-lapse movies were usually first normalized to create a constant overall intensity and compensate for the increasing intensity over time due to protein binding to the membrane.

#### Quantification of Bundle Width

To estimate the width of membrane-bound FtsZ bundles, fluorescence images were binarized using the adaptive threshold plugin for ImageJ. This plugin corrects for non-homogeneous background intensities, overcoming the limitation of conventional threshold methods. The background was removed using a local threshold corresponding to the mean intensity of an area of 20×20 pixels without any further subtraction. The binarized time lapse movie was then processed with the despeckle filter in ImageJ to remove small particles. Next, we calculated the Euclidean Distance Map (EDM) of every frame using the Distance Mapping function of ImageJ. This transformation results in a grey scale image, where the grey value of each pixel represents the shortest distance to the nearest pixel in the background. Accordingly, bundle widths correspond to the local peak intensities multiplied by 2.

The mean bundle width for every frame of the movie was calculated by identifying the peak intensities for each line and column of the image using a MATLAB script. This value could then be plotted as a function of time. The characteristic bundle width for different concentrations of ZapA and respective mutants was obtained by taking the time-average at steady state.

#### Architecture analysis

For a quantitative description of the filament architecture, we first calculated an orientation field of the pattern by calculating a gradient squared tensor at every position of a fluorescence micrograph^28,29^. This analysis was performed either using the OrientationJ plugin for ImageJ or using a custom python code based on the *scikit-image* package (Fig. 2A). For this method, a unit vector *u*_*θ*_ = (*cosθ*, *sinθ*) is assigned to all pixels in the image and their directional derivative is measured:

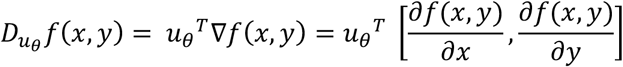

Where *f*(*x*, *y*) is the local grey value and ∇*f*(*x*, *y*) its corresponding gradient in x and y. The algorithm then finds the direction *u*_*φ*_, where the derivative is maximized over the region of interest (ROI):

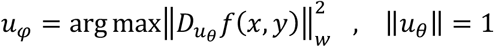

where *w*(*x*, *y*) ≥ 0 is related with the function that describes the ROI. Thus, the magnitude of each vector is proportional to the image contrast in a given direction and the average local direction is given by a weighted sum of all the vectors. A Gaussian filter, with a variance σ, governs the effective region of interest in which the orientation should be projected, i.e. the approximate dimension (in pixels) of the local structures in the image. A range of different σ was tested to find the best orientation field that could describe our data. This parameter was settled as 4 pixels for the analysis.

From a standard inner-product manipulation we get:

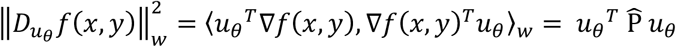

Where the first eigenvector of the so-called structure tensor matrix, 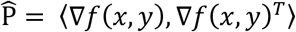 defines the local dominant orientation. The final output is a two-dimensional orientation field, *φ*(*x*, *y*), where each pixel has been replaced by unit vectors.

To estimate the curvature (*k*) of the orientation field, we used the definition from ref. ^28^, where it is described as the rate of change in the local orientation in the direction perpendicular to that orientation:

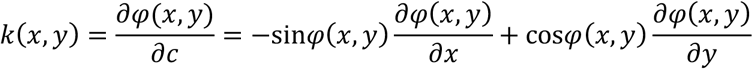

Where *c* corresponds to the axis perpendicular to the orientation. From this, we generated a color map containing *k* [*x*_*j*_ *y*_*i*_], which represents the curvature in *φ* at position x, y (Fig. 2B). Different colors (red and blue) indicate which way the curves bend (left or right). This depends on the way we define the parameters, i.e. which way we walk along the curve. Accordingly, as the direction of curvature is not a relevant parameter in our analysis, we plotted the distribution of absolute values for every experiment (mirrored negative values). The mean curvature for each image 〈*k*〉 was given by the mean half time of the mono-exponential decay fitted to these distributions (Fig. 2C).

To obtain a quantitative description of the spatial order of the filament network, we applied a nematic orientational correlation function, *S*(*r*), to the angles in the orientation field. This function compares the relative orientation of each angle with all the surrounding angles separated by an increasinsg distance *r*:

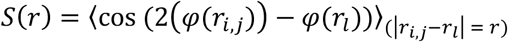

Where *φ*(*r*_*i,j*_) is the orientation angle at the position *r*_*i,j*_ and *φ*(*r*_*l*_) corresponds to all the surrounding angles at the position (*r*_l_), given that |*r*_*i,j*_ − *r*_*l*_| = *r*. The angular brackets indicate an average over the indicated range and the presence of the ‘2’ in the correlation function reflects the nematic symmetry of the system [−*π*/2, *π*/2]. As a result, we describe the average distance range over which the angles share a common orientation in each image. Each correlation curve typically goes from 1, where *r* is small and correlation is higher, to zero, where the correlation decreases drastically as the *r* increases. Accordingly, more ordered arrangements will display a higher spatial correlation at higher correlation lengths. To better visualize fluctuations in this parameter we identified a correlation length as *r* such that *S*(*r*) = 0.5 (intersection) as a function of time.

#### Image Autocorrelation

To quantify the reorganization dynamics of the membrane-bound FtsZ filament network, we performed a temporal correlation analysis using the <u>ImageCorrelationJ</u> plugin for ImageJ^30^. This method calculates the Pearson cross-correlation coefficient (PCC) between two frames with an increasing time lag between them (Δt)^31^. The mean intensities in local areas of 3×3 pixels were used to calculate the PCC between two different images, which was then plotted as a function of Δt (Fig.3B). For highly dynamic image pattern, the PCC typically decays rapidly with Δt, but slowly for persistent structures. The corresponding correlation curve was fitted to a mono-exponential decay, and the half-time (correlation time, *τ*) was used to compare a set of individual experiments.

#### Treadmilling Velocity Analysis

To quantify treadmilling dynamics, we have developed an automated image analysis based on differential image stack obtain from the ImageJ ImageCalculator command. First, two frames separated by 10s were subtracted from one another to generate a new time-lapse movie showing directionally moving fluorescent speckles. These speckles correspond to growth or shrinkage of filaments at a given position, (Fig. 4A and S3A). Next, the TrackMate^32^, a particle tracking toolbox available for ImageJ, was used to identify and track fluorescent speckles or filament treadmilling and to obtain detailed information regarding their displacement and velocity. Moving speckles were detected using the LoG (Laplacian Gaussian) detector with an estimated diameter of 1 μm. We used TrackMate’s quality criterion to select only the best 5% tracked particles and from these, we discarded particles with a signal-to-noise ration lower than 0.7. To build the final trajectories we used the “Simple LAP tracker” with a “Max Linking Distance” of 0.5μm, a “Maximal gap-closing distance” of 1μm and “Max frame Gap” of 2 frames, and we only considered for analysis trajectories longer than 6sec.

Reconstructed trajectories were further analyzed using a Matlab script based on @msdanalyzer^33^ toolbox to obtain information regarding the velocity and directionality of the moving speckles. The mean velocities obtained from TrackMate followed a normal distribution and were fitted to a Gaussian function to extract the mean velocity at each condition (Fig.4). In addition, we calculated the autocorrelation of the velocity vectors, i.e, angles of the normalized displacement vectors were compared pairwise as a function of an increasing time interval (Δt) and the correlation coefficient (V_corr_) was given by the cosine of the angle difference. This kind of analysis provided information about directionality since random motion particles tend to show velocity vectors completely uncorrelated (V_corr_ = 0 for all Δt) while particles with a directed migration display highly correlated velocity vectors (V_corr_ > for all Δt).

#### Single Molecule Tracking

Single molecule experiments were performed as described in ref. ^34^. Briefly, individual FtsZ monomers were resolved at single molecule level by mixing small amounts of Cy5-labelled FtsZ (less than 2nM) to a background of 1.5 μM Alexa488-labelled FtsZ. To track individual ZapA molecules, we followed a similar strategy by adding small amounts of ZapA-Cy5 to a background of unlabeled ZapA, always in the presence of Alexa488-FtsZ 1.5μM. Experiments were typically recorded using minimal laser power to visualize single molecules, with an exposure time of 50 ms and a varying acquisition time (114ms to 1s) for further photobleaching correction ^34^. Single molecules were identified and tracked using the automated particle-tracking platform TrackMate (https://imagej.net/TrackMate), with the following parameters: a particle size of 0.5 μm, particles localized for at least two frames were considered, linking max distance and gap closing distance of 0.5 μm, and 2 frames for the gap closing max. The mean lifetime of single molecules was obtained by fitting a mono-exponential decay to the lifetime distribution.

#### Fluorescence recover after photo bleaching (FRAP)

For FRAP experiments, we allowed the system to reach the steady state and a small area of the membrane was bleached using a high laser intensity. To obtain the half-time of the recovery, a Jython macro script for ImageJ (Image Processing School 8 Pilsen 2009) was used to fit the fluorescence recovery to *I*(*t*) = *a* (1 − exp (−*bt*)), where I(t) is the intensity value corrected for photobleached effects. FRAP experiments were acquired with an exposure time of 50msec and an acquisition time of one frame every 114msec.

#### Statistical analysis

For statistical analyses, two-tailed Student’s *t*-tests were performed using a python script. A *p*-value of < 0.05 was considered as statistically significant. For the boxplots, boxes are represented from 25th to 75th percentiles, the whiskers extend 1.5x the standard deviation and the mean value is plotted as a line in the middle of the box.

---

**Figure S1. Bundle width measurements of FtsZ/FtsA filament networks for ZapA WT and mutants**

**(A)** Snapshots of FtsZ pattern emerging from its interaction with FtsA and 0.5μM ZapA **(i)** or 1.5μM ZapA **(ii). (B)** Estimated 〈*δ*〉 over time for FtsZ/FtsA filament pattern in each condition (ZapA 0.5 μM, brown, n=5; 1.5Μm, dark green, n=7). **(C)** Snapshots of FtsZ pattern emerging from its interaction with FtsA and 6μM ZapA I83E **(i)** or 6μM ZapA R46A **(ii). (D)** Estimated 〈*δ*〉 over time for FtsZ/FtsA filament pattern in each condition (ZapA I83E, brown, n = 6; ZapA R46A, dark green, n = 4). Curves depict the mean and standard deviation (shade error bands) of independent experiments. **(E)** To estimate bundle width, each frame in the raw data **(i)** was binarized using a threshold plugin **(ii)** and then the Euclidean distance was calculated for every pixel in that frame **(iii).** Local peaks in the final gray scale image correspond to half of the bundle width (red line). **(F)** The mean bundle width over time was given by the peak of widths distribution of each frame without (left, black to blue) and with ZapA (right, black to red). Scale bars, 5μm.

**Figure S2. ZapA decreases FtsZ/FtsA filaments over time**

**(A)** Orientation fields and corresponding curvature maps for FtsZ filament network without (upper panel) or with 6.0 μM ZapA (lower panel) over time. Scale bars, 5 μm. **(B)** Mean curvature,〈*k*_*SS*_〉 and **(C)** Spatial correlation, 〈*S*(*r*)_*SS*_〉, at steady-state for FtsZ/FtsA filaments alone (blue) or upon addition of 6.0 μM ZapA (red). Curves depict the mean and standard deviation (shade error bands) of independent experiments (for〈*k*_*SS*_〉, FtsZ, n=9, ZapA, n=14; for 〈*S*(*r*)_*SS*_〉, FtsZ, n=11, ZapA, n=13).v

**Figure S3. Treadmilling speed of FtsZ/FtsA filaments**

**(A)** Illustration showing the outcome of the image subtraction procedure if the treadmilling was happening in parallel bundles (left panel) or in antiparallel filament bundles (right panel). For antiparallel bundles, we expect that speckles corresponding to growing and shrinking ends to initially overlap but then move in opposed directions. **(B)** Speckles intensity from several differential time-lapse movies (growth). Curves depict the mean and standard deviation (shade error bands) of independent experiments (FtsZ, n=5, ZapA, n=5). ZapA does not change the intensity of filament speckles, therefore it does not change the polarity of filaments within FtsZ bundles.

**TableS1. Statistical significance matrix**

Each matrix shows the *p value* for all the pairwise combinations of a two-tailed Student’s t-tests.

