## Supplementary Information for "ZapA stabilizes FtsZ filament bundles without slowing down treadmilling dynamics"

### Supplemental Figure S1

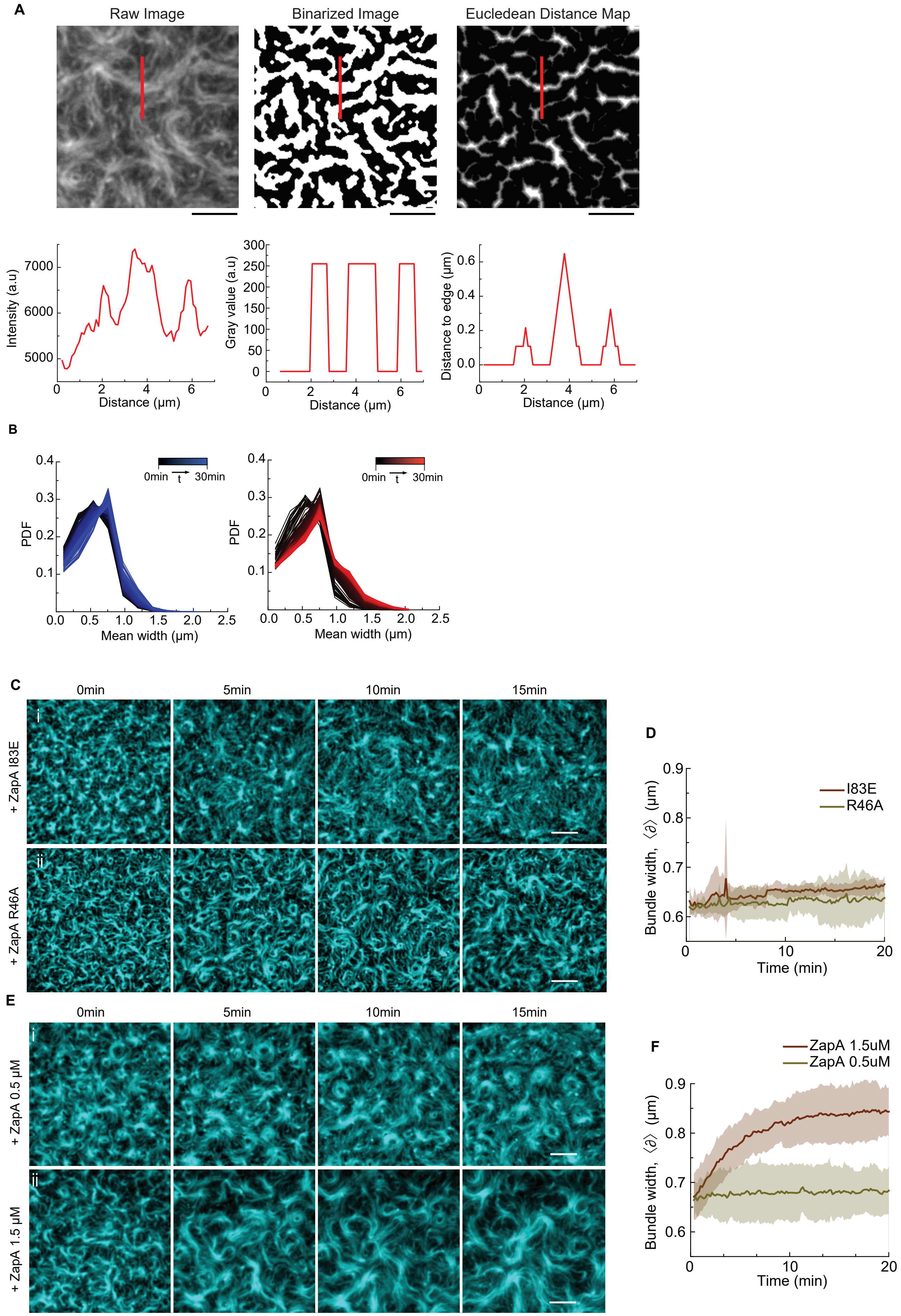

### Supplemental Figure S2

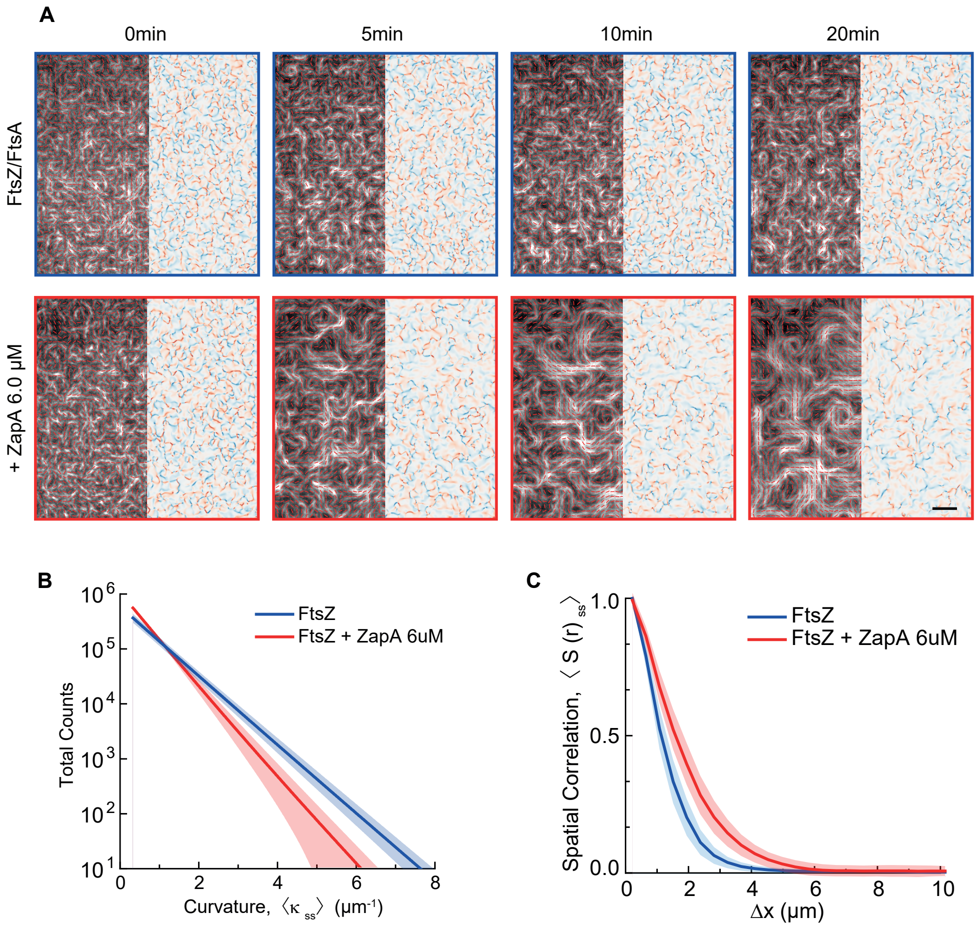

### Supplemental Figure S3

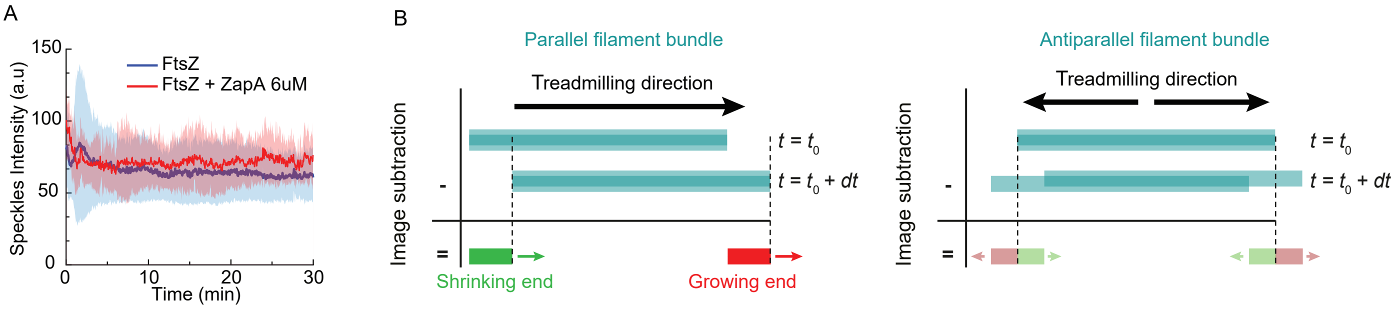

### Table S1: Statistical significance matrix

#### Bundle Width

| | FtsZ | ZapA wt | ZapA I83E | ZapA R46A |
|---|---|---|---|---|
| FtsZ | - | 6.5E-03 | 0.13 | 0.081 |
| ZapA wt | *** | - | 9.6E-06 | 2.61E-05 |
| ZapA I83E | n.s | *** | - | 0.47 |
| ZapA R46A | n.s | *** | n.s | - |

#### Curvature

| | FtsZ | ZapA wt | ZapA I83E | ZapA R46A |
|---|---|---|---|---|
| FtsZ | - | 5.8E-08 | 0.02 | 0.782 |
| ZapA wt | *** | - | 3.4E-04 | 3.14E-03 |
| ZapA I83E | * | *** | - | 0.02 |
| ZapA R46A | n.s | *** | * | - |

#### Correlation Length

| | FtsZ | ZapA wt | ZapA I83E | ZapA R46A |
|---|---|---|---|---|
| FtsZ | - | 1.6E-04 | 0.16 | 0.710 |
| ZapA wt | *** | - | 2.1E-03 | 1.10E-04 |
| ZapA I83E | n.s | *** | - | 0.13 |
| ZapA R46A | n.s | *** | n.s | - |

#### Correlation Time

| | FtsZ | ZapA wt | ZapA R46A | ZapA I83E |
|---|---|---|---|---|
| FtsZ | - | 2.0E-04 | 0.421 | 0.082 |
| ZapA wt | *** | - | 2.4E-03 | 0.044 |
| ZapA R46A | n.s | *** | - | 0.242 |
| ZapA I83E | n.s | * | n.s | - |

#### Directional Persistence

| | FtsZ | ZapA wt | ZapA I83E | ZapA R46A |
|---|---|---|---|---|
| FtsZ | - | 1.5E-04 | 0.450 | 0.760 |
| ZapA wt | *** | - | 0.030 | 3.2E-03 |
| ZapA I83E | n.s | * | - | 0.380 |
| ZapA R46A | n.s | *** | n.s | - |

#### SPT Lifetimes Measurements

| | FtsZ | FtsZ + ZapA WT | ZapA WT |
|---|---|---|---|
| FtsZ | - | 0.470 | 4.4E-07 |
| FtsZ + ZapA WT | n.s | - | 7.1E-04 |
| ZapA WT | *** | *** | - |

#### FRAP half-times

| | FtsZ | FtsZ + ZapA WT | ZapA WT |
|---|---|---|---|
| FtsZ | - | 0.140 | 1.4E-04 |
| FtsZ + ZapA WT | n.s | - | 1.7E-07 |
| ZapA WT | *** | *** | - |
